# Assessing Motivations and Barriers to Science Outreach within Academia: A Mixed-Methods Survey

**DOI:** 10.1101/2021.10.28.466319

**Authors:** Nicole C. Woitowich, Geoffrey C. Hunt, Lutfiyya N. Muhammed, Jeanne Garbarino

**Affiliations:** Department of Medical Social Sciences, Feinberg School of Medicine, Northwestern University, Chicago, Illinois 60611; American Society for Microbiology. Washington D.C. 20036; Division of Biostatistics, Department of Preventive Medicine, Feinberg School of Medicine, Northwestern University, Chicago Illinois 60611; RockEDU Science Outreach, The Rockefeller University, New York, NY 10065

## Abstract

The practice of science outreach is more necessary than ever. However, a disconnect exists between the stated goals for science outreach and its actual impact. In order to examine one potential source of this disconnect, we undertook a survey-based study to explore whether barriers to participation (either intrinsic or extrinsic) in science outreach exist within the academic community. We received responses to our survey from 530 individuals, the vast majority of whom engage in some type of science outreach activity on an annual basis. Those who engage in outreach report doing so for both personal and altruistic reasons, and having high (yet varied) levels of comfort with performing outreach activities. Respondents also report the existence of several significant yet surmountable barriers to participation, including lack of time and funding. Our findings demonstrate that both levels of participation in, and attitudes toward, science outreach within the academic community are generally favorable, suggesting that the general ineffectiveness of science outreach is due to other causes. We place our findings within the context of the broader science outreach, science communication and public engagement literature. We make recommendations on how existing approaches and infrastructure can, and must, be changed in order to improve the practice.

## Introduction

Never has the importance of effective science outreach been so well demonstrated as with the ongoing COVID-19 pandemic. As misinformation piles up and standard scientific processes get challenged on an increasingly frequent scale, there is an urgent need for cogent, lucid dialogue between the scientific and medical communities and broader publics. Yet the foundations upon which such engagements lie, such as trust, warmth and competence, are wobbly at best, leading to ineffective execution that has only furthered the divide between scientists and nonscientists (1). Before we as a scientific community can even begin to approach a cure, we must first diagnose the cause(s) of this disconnect.

Science outreach, as a general concept, is not new. Sometimes referred to as “science engagement” or “public engagement with science,” science outreach can be generally described as a framework that allows scientific communities from academic/institutional contexts to meaningfully connect with non-scientific communities around a collection of overlapping goals, which are met through the application of effective science communication and/or informal education practices, and are characterized by reciprocal learning for all involved (2,3). From natural philosophy lectures during the Renaissance era to the advent of science museums in the early 20th century to the use of social media channels in the 21st century, we have seen science outreach take many exciting, adaptable forms.

Unfortunately, within academia, science outreach has traditionally been perceived as a low-status task (4,5) that is typically performed by graduate students and early-career faculty, the majority of whom are women (4–7). In addition, significant barriers to participating in outreach exist, such as limited time and disapproval from peers or supervisors (5–7). We know from earlier work that institutions of science value research endeavors above all else, resulting in the marginalization of staff involved in science outreach efforts, as well as a scarcity of hard funding and infrastructure for such initiatives. Training for scientists wishing to contribute to these efforts is similarly lacking (8). The pursuit of science outreach and engagement activities is also often frowned upon by many members of the scientific community, whether because these activities are considered peripheral, or because weighing in on a politically charged subject is deemed “too risky” (9).

With so many variables affecting the implementation and execution of science outreach initiatives, it has been challenging to identify the most pertinent issues that need addressing. Though several social science research groups are working to understand the science of science outreach, a broad understanding of this field and how it fits within an evolving scientific ecosystem is limited. Often, reports on science outreach are centered on highly contextual circumstances, such as examining the role of a specific engagement program on science identity. While studies of this nature offer specific insights, these efforts are not relevant in understanding how to scale and sustain effective science outreach efforts associated with academic scientists and the institutions within which they conduct their research. Moreover, few peer-reviewed studies have examined science outreach’s relationship and utility to the scientific research enterprise. To date, only a small number of studies have focused on scientists’ involvement with science outreach (4,6,7,10–12). While informative, these studies have several limitations as they usually contain small sample sizes (4,6,7,10), are not specific to the U.S. research enterprise (5,11), or provide perspectives from a single scientific field (12).

As the nexus of science and society continues to mandate a shift in favor of science outreach and the practices of science communication and engagement, we must continue to explore the tensions that arise between academic science and these activities. Here, we contribute to the growing social science research surrounding the profession and practice of science outreach in order to better understand how the perceptions and motivations for participation in these activities has -- or has not -- shifted among academic STEM trainees and professionals. These data, in combination with existing findings, can help establish a baseline understanding of this rapidly growing field, which is an essential exercise if we are to sustain, or even expand, these efforts in ways that make genuine impact on our society.

## Methods

A survey instrument designed to assess scientists’ attitudes towards science outreach and motivations or barriers to participation was developed by the authors and piloted by a convenience sample of colleagues engaged in science outreach at varying levels of frequency. Based on the feedback from the pilot study, the survey instrument was further revised for clarity. The full survey instrument can be found in Supplementary File 1. A social media-based, snowball sampling strategy was utilized to distribute the survey (Qualtrics, www.qualtrics.com) between January and March 2020. Subsequently, the survey link received 1,796 unique clicks and was primarily accessed through email (72%), Twitter (32%), and other social media platforms (6%). Inclusion criteria included graduate students, postdoctoral fellows, faculty, and staff at academic institutions within the United States who self-identified as belonging to a science, technology, engineering or mathematics discipline.

### Quantitative Data Analyses

Demographics and survey respondents’ academic roles were summarized using frequencies and percentages. Categorical survey responses were reported using percentages. Median, mean, and standard error of the mean were computed for ordinal survey responses. Chi-square tests were used to compare participation in science outreach activities by academic role, academic rank, and gender. Differences in ordinal responses to questions related to comfort in science outreach participation were assessed using Kruskal-Wallis tests.

### Qualitative Data Analyses

Open ended survey comments were analyzed using a thematic approach. Two of the authors (NCW and JG) independently read and manually coded participant comments using deductive methods. Comments related to the statement, “I participate in science outreach because…” were assigned one of more of the following categories: Personal motivations; diversity, equity, inclusion and access; promoting education; societal relevance; inspiring the next generation; or other. Comments related to the statement, “I do not participate in science outreach because…” were assigned to one or more of the following categories: Time constraints; lack of professional incentive; unsure of opportunities; or other. Two of the authors (NCW and JG) discussed the comments to agreement.

The Rockefeller University institutional review board deemed this study exempt from further review. P values < 0.05 were considered significant.

## Results

### Characteristics of Survey Respondents

A survey instrument designed to assess scientists’ attitudes towards science outreach and identify motivations or barriers to participation in science outreach is described in detail in the Methods section. A total of 530 respondents completed the survey representing a range of scientific disciplines, career stages, and academic institutions. The majority of respondents were white (79%, *n*=416), women (72%, *n*=380), between the ages of 24 - 44 (66%, *n*=350) (Table 1). Respondents most frequently reported studying or working in New York (20%, *n*=104), Michigan (10%, *n*=53), Massachusetts (10%, *n*=53), California (7%, *n*=35), and Illinois (6%, *n*=31).

**Table 1.** Respondent Demographics

|  | N | % |
| --- | --- | --- |
| **Total** | 530 | *(100)* |
| **Gender** |  |  |
| Female | 380 | 72 |
| Male | 137 | 26 |
| Non-binary, Non-conforming, prefer to self-describe or not say | 13 | 2 |
| **Race** |  |  |
| White | 417 | 79 |
| Asian or Pacific Islander | 57 | 11 |
| Black or African American | 17 | 3 |
| Multiracial | 19 | 4 |
| Other, Prefer to self-describe or not say | 20 | 4 |
| **Ethnicity** |  |  |
| Hispanic or Latino/a/x | 49 | 9 |
| Non-Hispanic or Latino/a/x | 481 | 91 |
| **Age** |  |  |
| 18-24 years old | 46 | 9 |
| 25-34 years old | 233 | 44 |
| 35-44 years old | 117 | 22 |
| 45-54 years old | 58 | 11 |
| 55-64 years old | 53 | 10 |
| 65-74 years old | 19 | 4 |
| Prefer not to say | 4 | 1 |
| **US Location** |  |  |
| New York | 104 | 20 |
| Massachusetts | 53 | 10 |
| Michigan | 53 | 10 |
| California | 35 | 7 |
| Illinois | 31 | 6 |
| Other | 205 | 39 |
| Not disclosed | 49 | 9 |

Next, respondents were asked to describe their educational background, academic role and institution type (Table 2). The majority of respondents hailed from the biological sciences (57%, *n*=304) and had earned a PhD in their respective STEM discipline (54%, *n*=288). Respondents held varying roles in academia such as faculty (30%, *n*=159), graduate students (33%, *n*=177), postdoctoral fellows (15%, *n*=77), and staff / other (22%, *n*=117), and worked or studied at a research-intensive institution (74%, *n*=391). Fifty respondents (9%) indicated that they held a leadership role such as a Department Chair, Dean, Provost, or President. Of the faculty respondents (*n*=159), 67% held tenure track positions at the rank of assistant (32%, *n*=34), associate (30%, *n*=32), and full (38%, *n*=40) professor (Table 3). Faculty respondents reported that their research was primarily supported by institutional or departmental funds (48%, *n*=51) and National Science Foundation (27%, *n*=29) or National Institutes of Health (13%, *n*=14) grants (Table 3).

**Table 2.** Respondent Roles and Fields within Academia

|  | Total |  |
| --- | --- | --- |
|  | N = 530 | 100% |
| **Education Level** |  |  |
| BS or BA | 130 | 25 |
| MS or MA | 92 | 17 |
| PhD | 288 | 54 |
| Other, Prefer not to say | 20 | 4 |
| **Institution Type** |  |  |
| Research Intensive | 391 | 74 |
| Primarily Undergraduate | 65 | 12 |
| Religiously-Affiliated | 17 | 3 |
| Minority Serving Institution | 10 | 2 |
| Other or Unsure | 47 | 9 |
| **Role in Academia** | 530 | 100 |
| Faculty | 159 | 30 |
| Graduate student | 177 | 33 |
| Postdoctoral fellow | 77 | 15 |
| Staff | 94 | 18 |
| Other | 23 | 4 |
| **Leadership Role** |  |  |
| Chair, Dean, Provost, or President | 50 | 9 |
| **Field of Study** |  |  |
| Biological or Life Sciences | 304 | 57 |
| Social Sciences | 33 | 6 |
| Mathematics | 30 | 6 |
| Geosciences | 28 | 5 |
| Engineering | 25 | 5 |
| Chemistry | 25 | 5 |
| Physics | 25 | 5 |
| Astronomy | 20 | 4 |
| Other | 40 | 8 |

**Table 3.** Faculty Roles, Rank, and Primary Funding Source

|  | N | % |
| --- | --- | --- |
| **Faculty Track (n=159)** | 159 | 100 |
| Tenure track | 106 | 67 |
| Non-tenure track | 48 | 30 |
| Other | 5 | 3 |
| **Faculty Rank (n=106)** | 106 | 100 |
| Assistant Professor | 34 | 32 |
| Associate Professor | 32 | 30 |
| Professor | 40 | 38 |
| **Primary Funding Mechanism** | 106 | 100 |
| Institutional or Departmental | 51 | 48 |
| National Science Foundation | 29 | 27 |
| National Institutes of Health | 14 | 13 |
| Private Foundations or Philanthropic | 12 | 11 |

### Respondent Participation in Science Outreach

First, respondents indicated how often they participate in science outreach activities with varying degrees of frequency (Figure 1). In this survey, science outreach was defined as activities which, “broadly represent the different types of interactions between scientists and nonscientists…[sometimes] referred to as public engagement with science, informal science education, or science communication.” Frequency was defined by the following criteria: Never (I do not participate in science outreach), Rarely (1-2 times/year), Sometimes (2-3 times/year), Often (6+ times per year), Always (It is part of my job description / I consider myself a science outreach professional). The frequency of which respondents participated in outreach significantly differed by their role in academia [*X*^2^ (12, *n*=530) = 97.26,*p*<.0001]. Fifty percent of staff members engage in outreach “Always,” or “Often,” compared to 40% of graduate respondents, 38% of faculty, and 17% of postdoctoral fellows. Postdoctoral fellows comprised the largest group (58%, *n*=45) of respondents who “Rarely” or “Never” participate in science outreach (*n*=42) (Figure 1B). The frequency of faculty participation in science outreach did not differ by tenure track status [X^2^ (2, *n*=159) = 1.52, *p*=0.465] (Figure 1C) or rank [X^2^ (2, *n*=106) = 5.96, *p*=0.202] (Figure 1D). However, frequency of participation in science outreach varied by gender [X^2^ (2, *n*=517) = 8.63, *p*=0.013], as men were more likely to report that they did not participate in outreach, or did so rarely (45%, *n*=62) compared to women (31%, *n*=119, Figure 1E).

**Figure 1.**
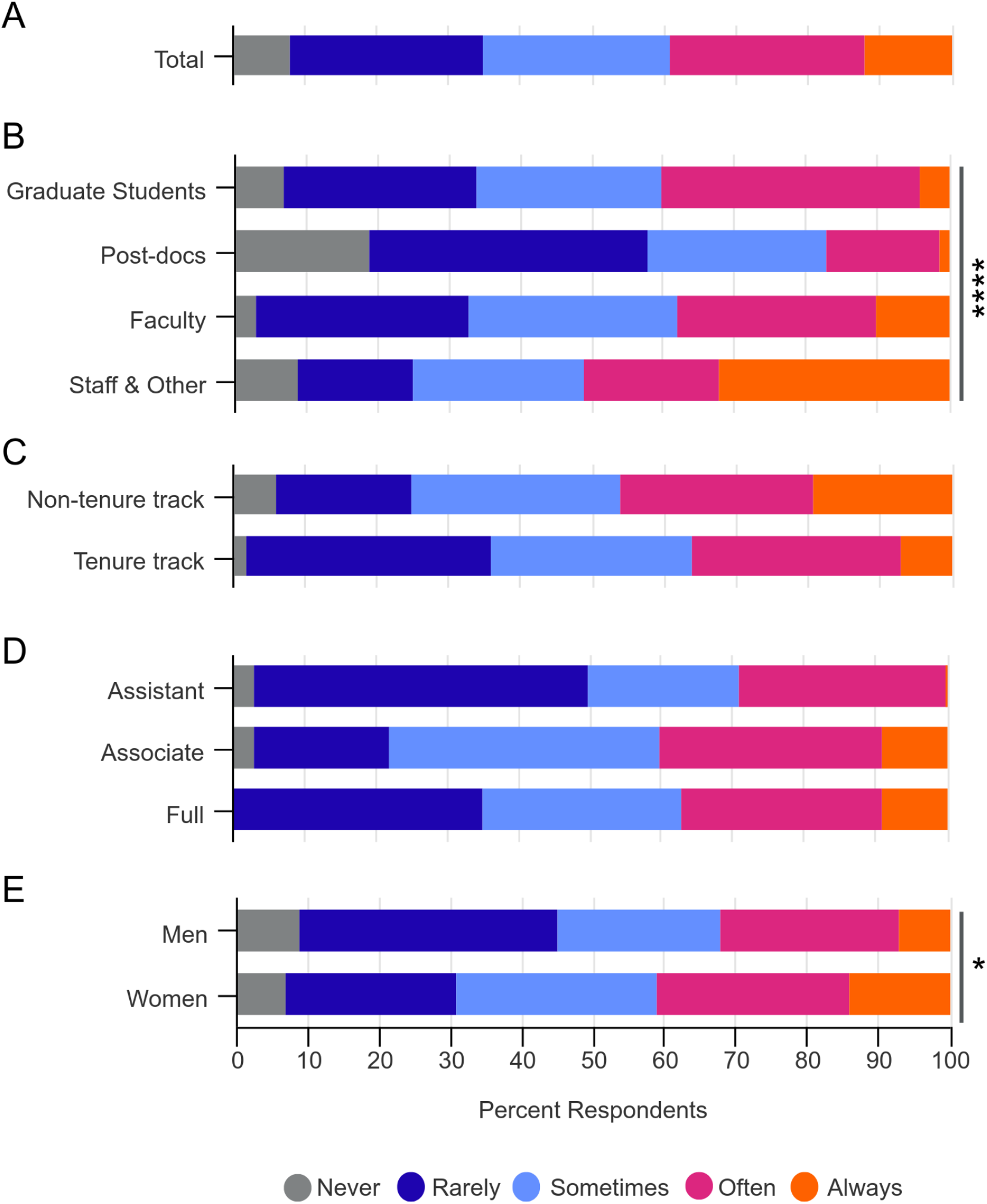
Comparison of respondents’ participation in science outreach activities. A) The majority of respondents (65%, n=344) participate in science outreach at least 3 or more times per year. B) Participation in science outreach varies by academic role *[X^2^* (12, *n*=530) = 97.26, *p*<.0001]. C) There were no differences in participation based on faculty track [X^2^ (2, *n*=159) = 1.52, *p*=0.465] or D) faculty rank [X^2^ (2, *n*=106) = 5.96, *p*=0.202]. E) Participation in science outreach differs by gender [X^2^ (2, *n*=517) = 8.63, *p*=0.013].

### Types of Outreach and Engagement

When asked about the types of science outreach activities that they participate in, the majority of respondents indicated that they engage with a K-12 student population (59%, *n*=305) or participate in public talks or lectures (54%, *n*=278) [Figure 2A]. In addition, many respondents also engaged in science outreach activities on social media (49%, *n*=251) or at science fairs (41%, *n*=214). Twenty-five percent of respondents (*n*=133) indicated that they have received some type of formal training in science outreach. Of the 75% of respondents who have not received any training (*n*=397), 50% (*n*=200) expressed an interest in doing so (Figure 2B).

**Figure 2.**
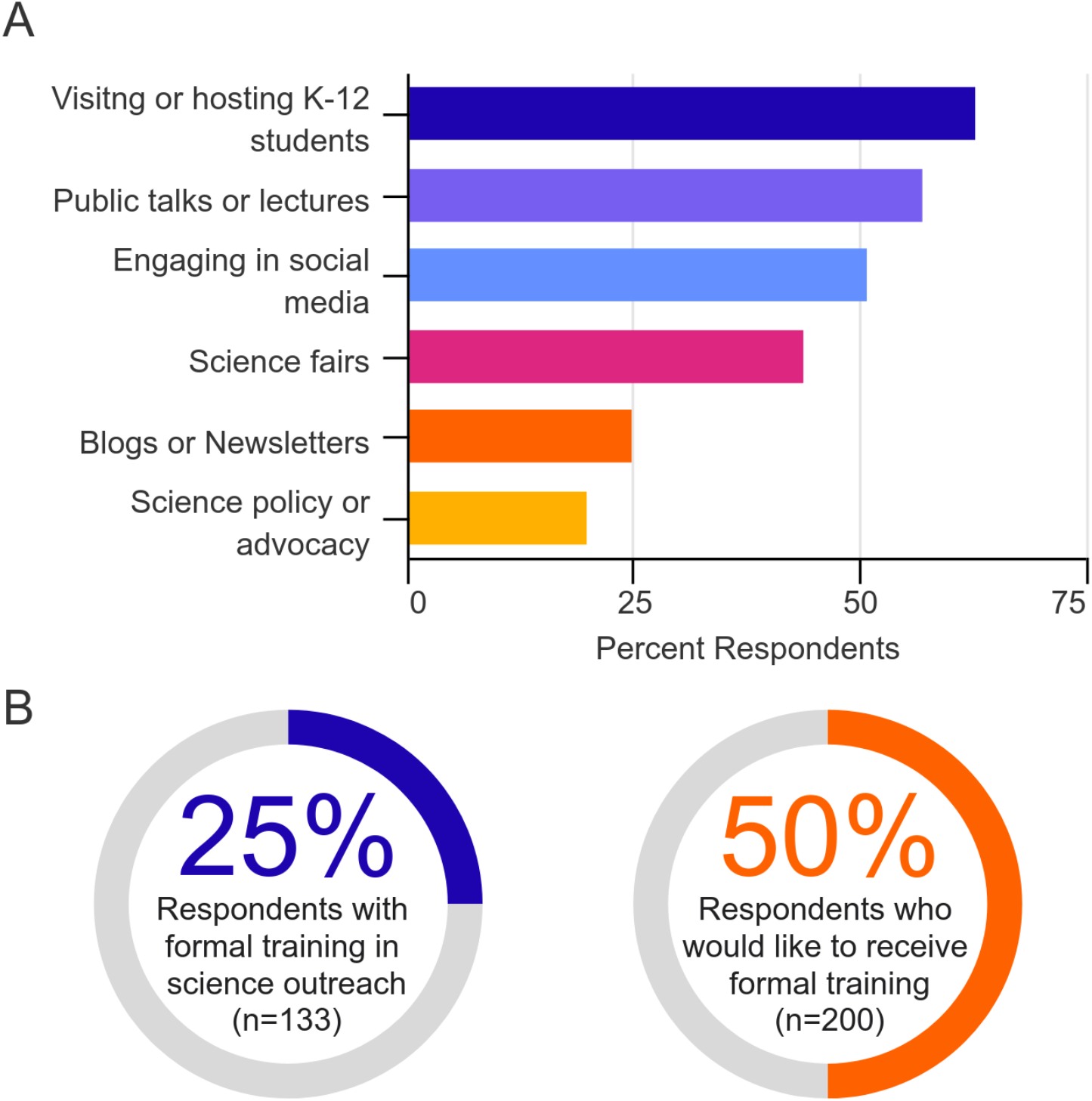
Respondents’ exposure to outreach activities and training opportunities. A) The percentage of respondents who engage in various science outreach activities. B) The percentage of respondents who have received or are interested in receiving formalized training in science outreach.

### Comfort Participating in Science Outreach

Respondents were asked how comfortable they would currently feel participating in a science outreach activity on a 10-point scale (1 being not comfortable at all, 10 being very comfortable) [Figure 3]. The median response was fairly high [Median (*Mdn)*=9, Mean (*M)*=8.3 Standard Error of the Mean (*SEM)*=0.09]. There was a significant difference in reported comfort levels among the various roles and ranks within academia [H=21.27, p=0.0003, *n*=530)] with faculty reporting the highest comfort levels (*Mdn*=9, *M*=8.8 *SEM*=0.13) compared to postdocs who reported the lowest comfort levels (*Mdn*=8, *M*=7.67 *SEM*=0.30) [Figure 3A]. There were significant differences in comfort levels based on respondents’ frequency participating in science outreach [*H*=137.5, p<0.0001, N=530], with increased comfort levels corresponding to greater frequency of participation [Figure 3B]. Those who did not participate in outreach had the lowest response in their comfort with outreach (*Mdn*=6.5, *M*=5.9 *SEM*=0.39) compared to those who always participated in outreach (*Mdn*=10, *M*=9.48 *SEM*=0.11). Respondent age was also a significant factor for who participates in outreach [*H*=15.1, *p*=0.019, *n*=530)], with those between the ages of 45-54 years old reporting the greatest comfort levels (*Mdn*=10, *M*=9.0 *SEM*=0.17) [Figure 3C]. There were no differences in comfort level in outreach participation based on gender [H=8.41, *p*=0.07, *n*=517] [Figure 3D].

**Figure 3.**
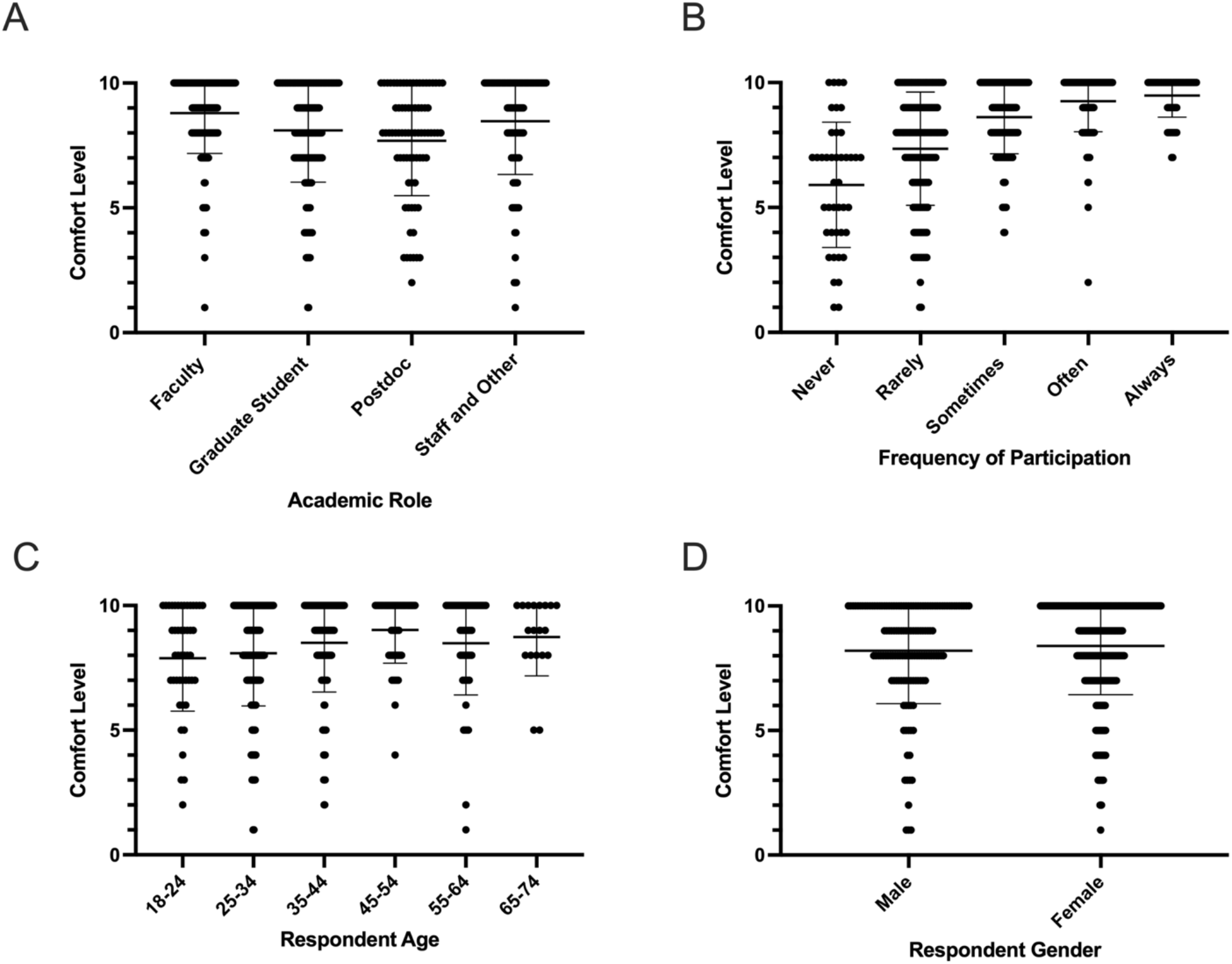
Comparison of respondents’ comfort level participating in science outreach activities. Respondents comfort level (1 – not being comfortable at all to 10 – being very comfortable) participating in science outreach significantly differed by by A) academic role [H=21.27, p=0.0003, n=530)], B) how often they participate in outreach levels [H=137.5, p<0.0001, N=530], and by C) age [H=15.1, *p*=0.019, *n*=530]. D) Respondents’ comfort level participating in outreach did not differ by gender [H=8.41, *p*=0.07, *n*=517].

### Perceived Utility of Science Outreach

Next, respondents were asked to select their level of agreement with statements pertaining to the utility of science outreach at the level of both the individual scientist and academic institution (Figure 4). The majority of respondents agreed that science outreach is a tool to establish relationships with their community (94%, *n*=498), establish interest in STEM careers (93%, *n*=492), and to educate or inform non-scientists about research findings (86%, *n*=455). The majority also agreed that science outreach was a tool for scientists to build transferable skills (67%, *n*=355). Only a minority of respondents agreed that science outreach was a tool to obtain research funding (34%, *n*=180). The majority of respondents also agreed that science outreach was a tool for academic institutions to establish relationships with their community (92%, *n*=486), create interest in STEM careers (92%, *n*=486), and educate or inform non-scientists about research findings (84%, *n*=443). The majority agreed that science outreach is a tool for academic institutions to address issues pertaining to diversity, equity, and inclusion (71%, *n*=378) and to promote cultural competency (63%, *n*=336). While 85% (n=452) of respondents agreed that science outreach is a tool to recruit students, only a smaller proportion of respondents indicated that it was a tool to recruit faculty members (34%, *n*=182). Lastly, the majority of respondents agreed that science outreach was a tool to attract philanthropic donors (67%, *n*=355).

**Figure 4.**
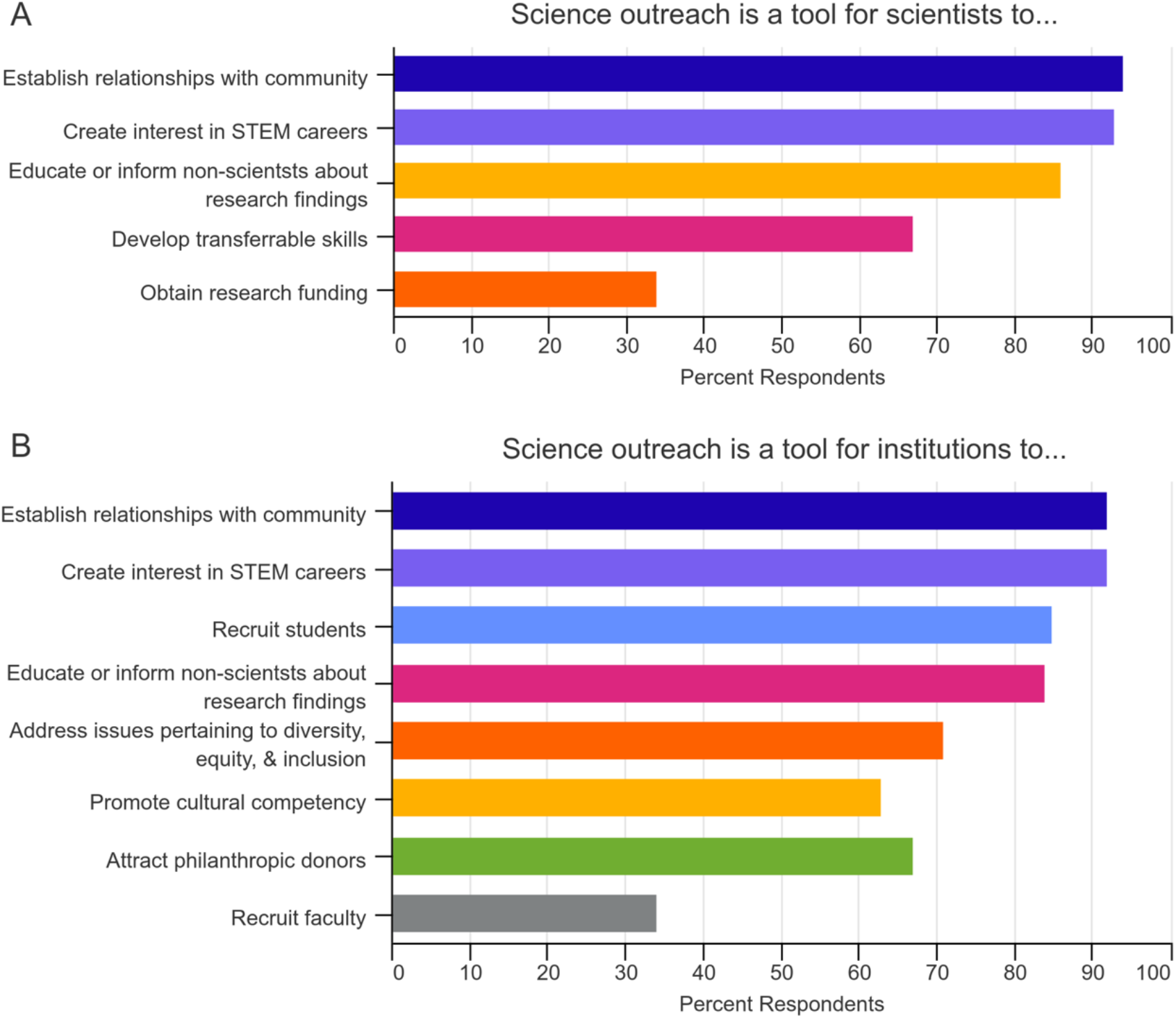
Respondents’ attitudes towards the perceived utility of science outreach. A) The percentage of respondents who agree that science outreach is tool for scientists to accomplish various tasks or initiatives. B) The percentage of respondents who agree that science outreach is a tool for academic institutions to accomplish various tasks or initiatives.

### Motivations & Barriers to Participation in Science Outreach

Respondents who indicated that they participated in science outreach with some level of frequency (either Rarely, Sometimes, Often or Always, *n*=488) were asked a series of questions surrounding their personal motivations to engage in outreach (Figure 5). Over 90% respondents indicated that they participate in science outreach because it is fun and enjoyable (92%, *n*=448). They also indicated that they participated in outreach to improve diversity, equity, inclusion, and access to STEM fields (91%, *n*=444) or to serve as a role model or mentor (86%, *n*=422). In addition, many respondents indicated that they participate in outreach to enhance their professional development (71%, *n*=345). Less than a quarter of respondents noted that grant funding requirements were a factor in their decision to participate in science outreach (23%, *n*=114).

**Figure 5.**
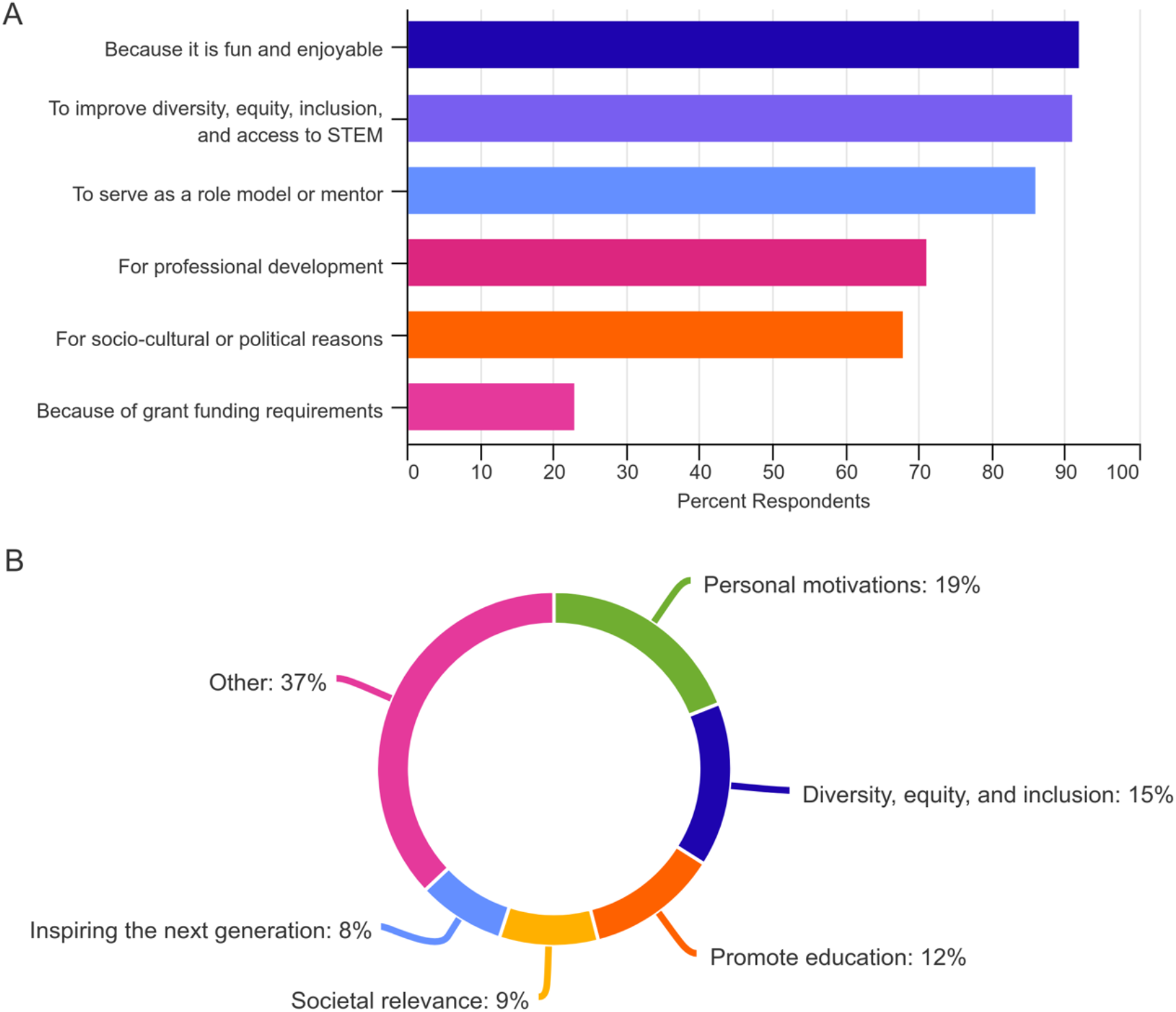
Respondents’ motivations for participating in science outreach. A) Percentage of respondents (n=488) who indicated that they participate in science outreach for various reasons. B) Percentage of codes (n=388) associated with respondents’ qualitative responses to the question, “I participate in science outreach because…”

Respondents had the option to complete the following free-response statement: “I participate in science outreach because…” Three-hundred and thirteen statements were independently reviewed and coded by two of the authors (NCW and JG) and discussed to agreement. Fifteen codes emerged, with a total of 388 occurrences. The proportion of codes assigned were: Personal motivations (20%, *n*=77), diversity, equity, inclusion and access (DEIA, 16%, *n*=62), promoting education (12%, *n*=48), societal relevance (10%, *n*=38), and inspiring the next generation (8%, *n*=32; Figure 5B). Respondents’ personal motivations focused on the direct benefit to the individual. For example, respondents indicated that they participate in science outreach because, “I value the experience and interactions…,” or “It gives my life meaning!” Other comments focused on the importance of improving diversity, equity, inclusion, and access to STEM fields. One respondent shared, “I believe it is essential in breaking down structures of privilege that influence who enters academia and sees themselves in science or not.” Another respondent noted, “I wouldn’t be here without [science outreach] and everyone should have access, support, and resources for equitable opportunity in science.”

A separate theme focused on science outreach as a tool to advance science education. One respondent noted that they participate in outreach, “To help non-scientists better understand scientific concepts.” Several other comments focused specifically on enhancing scientific literacy through outreach. “I’m passionate about improving science literacy in non-scientist members of the public,” shared one respondent. Another stated that: “Scientific literacy and critical thinking are important skills for the general public to have.” In addition to education, the societal relevance of science was another key motivator to engage in science outreach. Several respondents shared that, “It helps non-scientists understand why science matters to society,” and “It helps the taxpaying public understand what we do and, hopefully, to value it.” Lastly, many respondents indicated that they participated in science outreach to inspire the next generation of scientists. “It lets me show kids the opportunities they have in science (which I hadn’t been aware of as a kid),” shared a respondent. A different respondent echoed that sentiment, “Seeing others like myself in science helped allow me to imagine myself as a scientist, thus opening a door to a career pathway (and a way of advancing knowledge) I hadn’t previously considered.” Comments which were ultimately categorized as “Other” included 10 themes ranging in frequency from 0.5 - 7%. The response varied and discussed topics such as professional obligations, a sense of duty, obligation to taxpayers, and to build trust and/or combat misinformation.

Respondents also highlighted the major barriers to participating in science outreach (Figure 6A). Lack of time was reported as the most common barrier experienced by respondents (84%, *n*=412), followed by lack of funding (49%, *n*=238), whereas public disinterest in science (14%, *n*=67) and communication or language barriers (7%, *n*=32) were reported less frequently. Of the respondents who never participate in science outreach (*n*=42), the majority cited lack of time (67%, *n*=28) and lack of funding (69%, *n*=29) as major barriers to participation.

**Figure 6.**
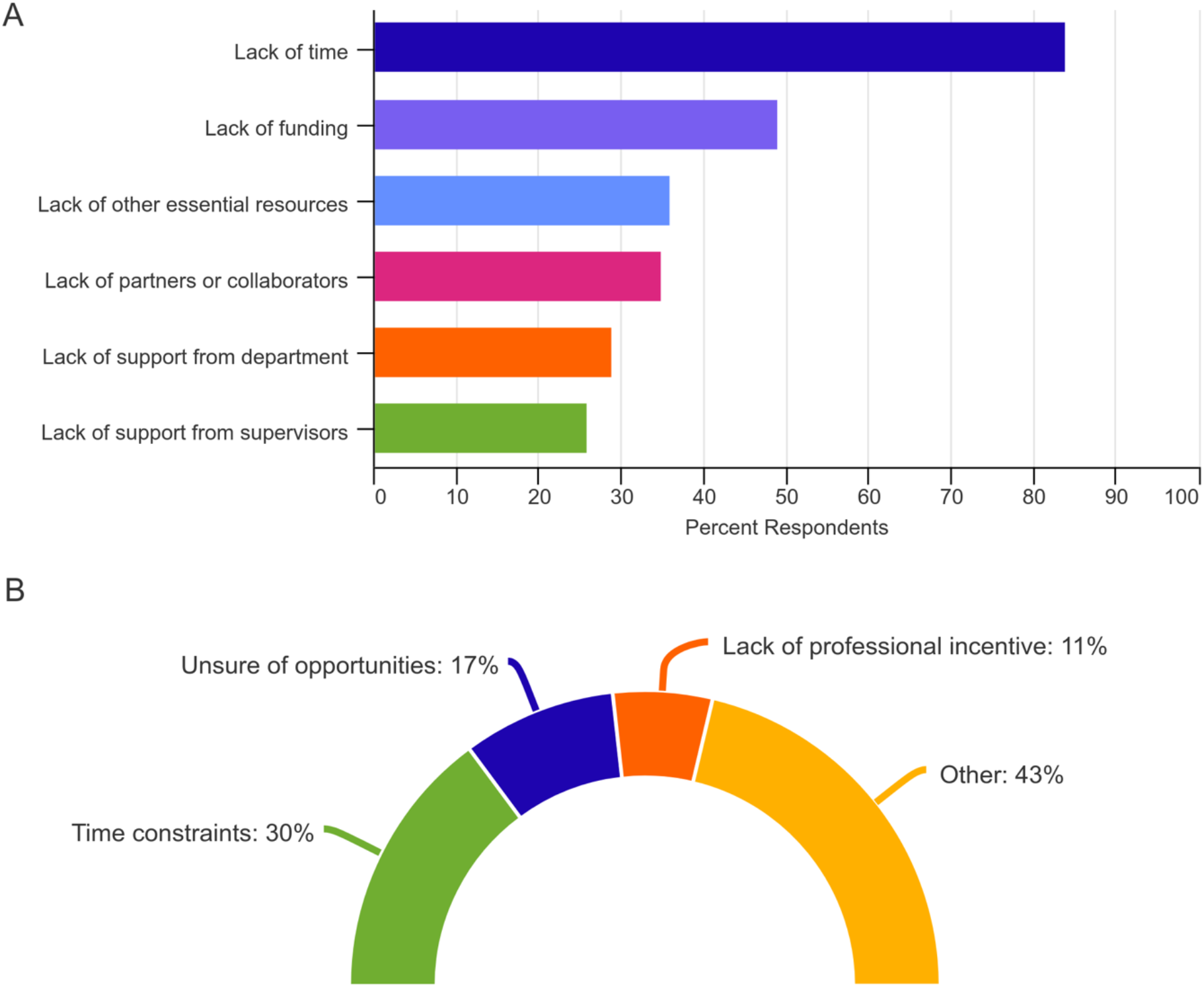
Barriers to respondents’ participation in science outreach. A) Percentage of respondents (n=488) who indicated that they experience barriers which hinder their participation in science outreach. B) Percentage of codes (n=31) associated with respondents’ qualitative responses to the question, “I do not participate in science outreach because…”

Respondents who indicated that they never participated in science outreach (*n*=42) were asked to complete the following free-response question: “I do not participate in science outreach because…” Thirty-one responses were coded by two of the authors (NCW and JG). Four codes emerged with a total of 47 instances (Figure 6B). Respondents cited time constraints as the top barrier to participating in outreach (30%, *n*=14). One respondent noted that, “I’ve been too busy working in the lab to participate in many other activities, and science outreach isn’t my number 1 priority for what to devote extra time to.” Another stated that, “There is already so much other work that needs to be done. Science outreach would just be one more thing to do without much [return on investment] for already stretched faculty and staff.” An additional major barrier was being unaware of opportunities to engage in outreach (17%, *n*=8). “I have not been able to easily find opportunities that fit my schedule/available time, and that are easy to get to participate in, and that are within my comfort zone,” shared one responded. Lack of professional incentive was another concern shared by several respondents (11%, *n*=5). As one respondent described, “It is not valued by the scientific community. That means it will not help me get a job after my postdoc. In such a hypercompetitive environment, how can I justify spending time on outreach when so much else gets piled on?” The remainder of the comments were categorized as “Other” and touched upon themes related to self-doubt (“I am shy and nervous about [participating]”), supervisor disapproval, (“I think my advisor would want me to do more research instead of participating in outreach,”) and professional limitations (“It is not an available option with my position,”).

## Discussion

The final sentence of the seminal 1985 publication, “The public understanding of science: report of the Royal Society’s ad hoc group” (colloquially referred to as the “Bodmer report”) issued one of the first broad clarion calls to the scientific community with respect to public engagement with science: “[O]ur most direct and urgent message must be to the scientists themselves: Learn to communicate with the public, be willing to do so and consider it your duty to do so.” (13). Thirty-six years later, where are we with regard to the perception of science outreach among members of academic STEM communities?

As shown by our results, it is encouraging to see that significant progress has been made towards answering the challenge issued by Bodmer, and many colleagues since. The members of the scientific community who we surveyed have an overwhelmingly positive view of science outreach, have received or would like to receive training, participate in large numbers in a variety of activities, and have high levels of comfort with their participation in outreach. Even more encouraging are the findings that participation in science outreach is overwhelmingly driven by an interest to showcase that science can be fun and full of wonder, and can help promote diversity, equity, inclusion, and access (DEIA) best practices -- issues that also resonate with many target audiences for science outreach activities. In addition, there has been widespread acknowledgement that the “public” is not a monolith, but is rather a collection of many groups of connected individuals, in which scientists, too, belong (14).

While the intentions driving these trends all appear to be moving in a positive direction with respect to attitudes towards and participation in public engagement with science, a larger question remains: has all of this energy, effort, expenditure of resources and change in mindset moved the needle at all in terms of what science outreach is actually trying to accomplish?

If we look beyond internal metrics and focus on external impact, we cannot confidently answer this question with a “yes.” Though many scientists have a positive outlook towards science outreach and believe that outreach is worthwhile (4,11,12,15), research has shown that the types of science outreach activities that are popular with the scientific community and are meant to democratize science (such as science fairs and festivals, public lectures, and K-12 classroom visits) are instead reaching limited audiences (16,17) and having minimal impact on perceptions towards, and understanding of, science (18).

Furthermore, even though decades of social science research have prompted an evolution in the description, cultivation and maintenance of relationships that scientists can develop with a variety of communities via science outreach (8), much of the qualitative feedback we received from respondents as to why they participate in science outreach generally suggests a deficit-model, top-down approach to science communication and engagement. This counterproductive approach, in which the representation of STEM to non-expert audiences is dictated by the dominant cultural norms of STEM, serves to attract only those who are already familiar with these cultural norms, and perpetuates much of the exclusive nature of science and science outreach, particularly to marginalized communities who, throughout history, have been discouraged and actively prevented from engaging in scientific processes (14,19).

When considered collectively, we should acknowledge that current science outreach efforts are collectively missing the mark, despite the significant uptick in individual scientists and laboratories attempting to create sustainable engagement efforts. It is unsurprising that among non-scientist audiences, science is not regularly seen as being relevant to the everyday issues at the forefront of public discourse (20). Additionally, even though our study suggests that motivations for participation in science outreach are often centered on improving issues related to DEIA in STEM, it is well-appreciated that the makeup of those who work or study in STEM fields does not reflect the demographics of the American population (21).

Additionally, we must consider the value of science outreach and engagement training for scientists. How do we reconcile the need to prepare scientists for engaging effectively with their communities, focusing on social justice issues to improve DEIA (8), with the notion that rigorous training may not actually be working to adequately prepare scientists for this work (22)? Perhaps the answer requires a more holistic approach that takes into consideration not just goals and outcomes we set for engagement, but also participatory motivations for all stakeholders --scientists and non-scientists alike. Through this study, we confirmed that many scientists are motivated to participate in outreach because they view science as fun and full of wonder, and have a desire to share these potentially powerful emotions with others. While it often feels taboo to make space for emotion in scientific environments, prompting “fun” emotions in science can actually enhance the likelihood that non-scientists will engage with science again in the future (23,24). This is particularly relevant when science outreach efforts are intended to serve minoritized communities (25).

We recognize that the findings from this study may be limited by our relatively small, yet not insignificant, sample size (N=530). However, the outcomes of this research are in sync with the handful of similar studies from the past 20 years that have examined scientists’ participation in, and attitudes towards, science outreach. Our findings suggesting that members of STEM communities hold highly positive attitudes towards outreach is echoed by studies from others (10,12,26). Similarly, our results demonstrate that fun and skill development are strong motivators for participating in outreach and align with results by Andrews and colleagues (6). Our study is also consistent with the outcomes of work performed by Ecklund et al and Johnson et al, who reported that women participate more than men in outreach (4,7). White women were highly represented in our respondent pool compared to other groups, which may reflect a demographic trend for the field of science outreach (27). It is worth acknowledging that both the PI and Co-PI for this study are white women. We posit that the overrepresentation in responses from this group may be, to some extent, a reflection of homogeneity in the authors’ academic and social networks -- a situation known as homophily that is present across industries, including economics and science (28,29).

A strength of our work, in contrast to previous studies, is that we were able to gather responses from individuals from all different career stages and academic fields, which may provide a more comprehensive overview of how science outreach can be incorporated across academic contexts. Looking at constraints on participation in outreach, Andrews et al and Ecklund et al report that a lack of time is one of the biggest barriers, though surprisingly the report from Poliakaff et al did not find time to be a barrier (6,7,10). As with our study, Ecklund et al, Rose et al, Andrews et al. all found that respondents felt outreach was not seen as being valued professionally (6,7,26). Our study found that 50% of respondents would like to receive formal training in science outreach, confirming conclusions from Poliakaff et al that providing training in this space would boost enthusiasm and overcome a major barrier to participation in science outreach activities (10). However, it is not as easy as just implementing some sort of training regimen -- as mentioned above, we see uneven results even with rigorous engagement for scientists.

Collectively addressing these outstanding issues requires yet another cultural shift, one that is less about motivating academic scientists to participate in outreach activities, and more about the design and implementation of those activities in context. As demonstrated by current crises, ranging from the COVID-19 pandemic to climate change to racial injustice, the scientific community writ large is still taking a deficit-model approach to communicating science in which audiences are passive recipients of information delivered using formal, top-down methods and outlets. It can be no surprise, consequently, that this approach has led to diminishing respect towards, and trust in, in science and the authority of scientists (30).

We cannot continue simply promoting the benefits of outreach and engagement within the scientific community. Instead, our efforts must align with those calling for inclusive science communication and engagement, an approach that demands more focus and attention be paid towards community-centric, bottom-up approaches to public outreach that tear down silos preventing collaboration and communication across various sectors and groups (9). This is especially true for universities, who are often at the industrial center of their respective geographic communities.

Clearly there is demand for science outreach training and opportunities, a welcomed trend that we hope continues to accelerate. Yet as we look forward, we argue that these offerings must be properly designed and distributed, in a manner that takes into account the best principles and practices of community engagement. To do so, we must prioritize closing the gap between research on science communication and public outreach and implementation of best practices. Furthermore, there is a need for stakeholder collaboration on developing ways to examine whether the trainings, activities and programs being implemented are actually achieving their stated aims.

Only by embracing this mindset of inclusivity and truly engaging with individuals and communities can the practitioners of science outreach achieve the lofty goals of improving DEIA, creating interest in STEM careers, and increasing awareness, understanding, and involvement with science. Despite improvements that have been made, none of us should be satisfied with the current state of affairs with regard to how science is practiced, taught and disseminated. We can, and must, do better.

## Supporting information

Survey Questions

## Author Contributions

Nicole C. Woitowich: Conceptualization, Resources, Software, Formal analysis, Supervision, Funding acquisition, Investigation, Methodology, Writing - original draft, Writing - review and editing

Geoff C. Hunt: Formal analysis, Writing - original draft, Writing - review and editing

Lutfiyya N. Muhammed: Formal analysis, Methodology, Writing - review and editing

Jeanne Garbarino: Conceptualization, Resources, Software, Formal analysis, Supervision, Funding acquisition, Investigation, Methodology, Writing - original draft, Writing - review and editing

## Acknowledgement

The authors thank the following individuals who participated in the pilot survey. In addition, they thank all of the individuals who participated in this study.

## Ethics

This work was reviewed and deemed exempt by the institutional review boards of The Rockefeller University and Northwestern University.

