## Supplementary material for "Assessing Motivations and Barriers to Science Outreach within Academia: A Mixed-Methods Survey": Survey Questions

**Title of Research Study: Integration of Science Outreach into the Academic Research Enterprise: Data-Driven Perspectives**

**Investigators:**
Nicole C. Woitowich, PhD
Northwestern University


Jeanne Garbarino, PhD
Rockefeller University


**Supported By:** National Science Foundation Grant no. 1854018 to Jeanne Garbarino, PhD and Nicole Woitowich, PhD

**Key Information:** The purpose of this study is to examine how science outreach currently exists within academic research institutions at the national level. You will be asked to complete an online survey which will take approximately 15 minutes to complete. There are no risks or benefits associated with taking this survey. All information collected is completely anonymous.

**Why am I being asked to take part in this research study?** You are being asked to take part in this research study because you are a graduate student, postdoctoral researcher, faculty, administration, or staff at a college, university, or research institution.

**What should I know about a research study?** Participation is voluntary, and you may discontinue participation at any time by closing your browser or leaving this webpage.

**What happens to the information collected for the research?** This survey is being hosted by Qualtrics and involves a secure connection. Terms of service addressing confidentiality may be viewed at http://www.qualtrics.com/research-suite/. No personally identifiable information will be collected.

**What else do I need to know?**
You will not receive compensation of any kind for participating in this study.

**Who can I talk to?** If you have questions, concerns, or complaints talk to the Principal Investigators, Dr. Nicole C. Woitowich at 312-503-1385 or Dr. Jeanne Garbarino at 212-327-7418. If you have any concerns about your experience while taking part in this research study, you may contact The Rockefeller University Institutional Review Board (IRB) at 212-327-8408. You can also speak with members of the Northwestern University IRB at 312-503-9338.

**Consent**
If you want a copy of this consent for your records, you can print it from the screen.
**By completing this survey, you are consenting to participate in this study.**

End of Block: Consent Form

Start of Block: Demographics

Do you study or work in a science, technology, engineering or math (STEM) field?

- ○ Yes
- ○ No

*Skip To: End of Survey If Do you study or work in a science, technology, engineering or math (STEM) field? = No*

Do you study or work at a 2- or 4-year college, university, or academic research institution within the United States?

- ○ Yes
- ○ No

*Skip To: End of Survey If Do you study or work at a 2- or 4-year college, university, or academic research institution with... = No*

What is your primary field of study?

- ○ Astronomy
- ○ Biological or Life Sciences
- ○ Chemistry
- ○ Computer Science and Technology
- ○ Engineering
- ○ Mathematics
- ○ Physics
- ○ Geosciences
- ○ Social Sciences
- ○ Other

Please select the option which best describes your college, university, or research institution:

- ◯ Community or Junior College
- ◯ Primarily Undergraduate College or University
- ◯ Research Intensive College or University
- ◯ Historically Black College or University
- ◯ Tribal College or University
- ◯ Religiously-affiliated College or University
- ◯ Minority-Serving Institution
- ◯ Other
- ◯ I'm not sure

Page Break

Would you be willing to provide the zip code of where your campus is located?

- ◯ No

*Invalid Logic Click Here to Edit Logic*

- ◯ Yes ________________________________________________

Page Break

What is your current gender identity?

- ◯ Male
- ◯ Female
- ◯ Gender non-binary or gender non-conforming
- ◯ Prefer not to say
- ◯ Prefer to self-describe

Which group most accurately describes your race?

- ◯ American Indian or Alaska Native
- ◯ Asian or Pacific Islander
- ◯ Black or African American
- ◯ White
- ◯ Multiracial
- ◯ Prefer to self-describe
- ◯ Prefer not to say

Which group most accurately describes your ethnicity?

- ◯ Hispanic or Latino/a/x
- ◯ Not Hispanic or Latino/a/x

Which of the following best describes your sexual orientation?

- ◯ Heterosexual
- ◯ Lesbian
- ◯ Gay
- ◯ Bisexual
- ◯ Asexual
- ◯ Pansexual
- ◯ Prefer to self-describe
- ◯ Prefer not to say

---

Please select your age range

- ◯ 18 - 24 years old
- ◯ 25 - 34 years old
- ◯ 35- 44 years old
- ◯ 45 - 54 years old
- ◯ 55 - 64 years old
- ◯ 65 - 74 years old
- ◯ 75 years or older
- ◯ Prefer not to say

---

Page Break

Select the highest degree you have obtained:

- ◯ BS or BA
- ◯ MS or MA
- ◯ PhD
- ◯ MD
- ◯ Other professional degree
- ◯ Prefer not to say

What is your current role in academia?

- ◯ Graduate Student
- ◯ Postdoctoral Fellow
- ◯ Faculty
- ◯ Staff
- ◯ Other ________________________________________________

*Display This Question:*

*If What is your current role in academia? = Faculty*

Please select your faculty track:

- ◯ Tenure track
- ◯ Non-tenure track (ex. Research faculty, Clinical Instructors, Lecturers)
- ◯ Other

*Display This Question:*

*If Please select your faculty track: = Tenure track*

Please select your academic rank:

○ Assistant Professor

○ Associate Professor

○ Professor

Do you hold a leadership position within academia? Examples include: Department Chair, Dean, Provost, or President

○ Yes

○ No

○ Unsure

**End of Block: Demographics**

**Start of Block: Science Outreach Experience**

For the next series of questions, we use the term “science outreach” to broadly represent the different types of interactions between scientists and non-scientists. You might refer to this as public engagement with science, informal science education, or science communication.

On a scale of 1-10, how comfortable would you be participating in a science outreach activity today?

| | 1 | 2 | 3 | 4 | 5 | 6 | 7 | 8 | 9 | 10 |
|---|---|---|---|---|---|---|---|---|---|---|
| Level of comfort (1 not at all comfortable - 10 being very comfortable) | | | | | | | | | | |

How often do you participate in science outreach activities?

- ○ I do not participate in science outreach activities
- ○ Rarely (1 - 2 times a year)
- ○ Sometimes (3 -5 times a year)
- ○ Often (6+ times a year)
- ○ Always - It is a part of my job description / I consider myself a science outreach professional

*Display This Question:*

*If How often do you participate in science outreach activities? != I do not participate in science outreach activities*

What types of science outreach activities do you participate in? Check all that apply:

- ☐ Visiting or Hosting K-12 Students
- ☐ Science fairs
- ☐ Public talks or lectures
- ☐ Podcasts
- ☐ Blogs or Newsletters
- ☐ Social media (Twitter, Facebook, Instagram)
- ☐ Website or Wiki Curation
- ☐ Science policy or advocacy
- ☐ Other

Have you ever received any formal training in science outreach?

- ◯ Yes
- ◯ No, I have not
- ◯ No, but I would like to
- ◯ Unsure

Page Break

In your opinion: Who *most often* participates in science outreach at your college, university, or institution? Check all that apply.

- ☐ Undergraduate Students
- ☐ Graduate Students
- ☐ Postdoctoral Fellows
- ☐ Faculty Members
- ☐ Staff
- ☐ Other ________________________________________________

Page Break

In your opinion: Who *most often* participates in science outreach at your college, university, or institution? Check all that apply.

- ☐ Men
- ☐ Women
- ☐ Gender non-binary or non-conforming individuals
- ☐ White (non-hispanic) people
- ☐ Persons of color
- ☐ LGBTQIA+ identifying
- ☐ People under the age of 40
- ☐ People over the age of 40
- ☐ Native english speakers
- ☐ Non-native english speakers
- ☐ Other ________________________________________________

Page Break

End of Block: Science Outreach Experience

Start of Block: Science Outreach Value

Please select your agreement with the following statement: *Science outreach is a tool for* ***scientists*** *to...*

| | Strongly Disagree | Disagree | Neutral | Agree | Strongly agree |
|---|---|---|---|---|---|
| Educate or inform non-scientists about research findings | ○ | ○ | ○ | ○ | ○ |
| Establish relationships with their community | ○ | ○ | ○ | ○ | ○ |
| Create interest in STEM careers | ○ | ○ | ○ | ○ | ○ |
| Develop transferrable skills | ○ | ○ | ○ | ○ | ○ |
| Obtain research funding | ○ | ○ | ○ | ○ | ○ |
| Maintain research funding | ○ | ○ | ○ | ○ | ○ |

Please select your agreement with the following statement: *Science outreach is a tool for **colleges, universities, or institutions** to...*

| | Strongly Disagree | Disagree | Neutral | Agree | Strongly agree |
|---|---|---|---|---|---|
| Educate or inform non-scientists about research findings | ○ | ○ | ○ | ○ | ○ |
| Create interest in STEM careers | ○ | ○ | ○ | ○ | ○ |
| Recruit students | ○ | ○ | ○ | ○ | ○ |
| Recruit faculty members | ○ | ○ | ○ | ○ | ○ |
| Address issues pertaining to diversity and inclusion | ○ | ○ | ○ | ○ | ○ |
| Promote cultural competency | ○ | ○ | ○ | ○ | ○ |
| Establish relationships with their community | ○ | ○ | ○ | ○ | ○ |
| Attract philanthropic donors | ○ | ○ | ○ | ○ | ○ |

Page Break

Please select the best answer for each of the statements below:

| | Yes | No | Unsure | Not Applicable |
|---|---|---|---|---|
| Science outreach is valued by my peers | ○ | ○ | ○ | ○ |
| Science outreach is valued by my supervisors | ○ | ○ | ○ | ○ |
| Science outreach is valued within my department | ○ | ○ | ○ | ○ |
| Science outreach is valued by my institution | ○ | ○ | ○ | ○ |
| Science outreach is valued by the scientific community | ○ | ○ | ○ | ○ |

---

On a scale of 1 - 10, how much do you personally value science outreach?

| | 1 2 3 4 5 6 7 8 9 10 |
|---|---|
| (1 - I do not value science outreach at all, 10 - I completely value science outreach) | |

End of Block: Science Outreach Value

Start of Block: Science Outreach Motivations / Limitations

*Display This Question:*

*If How often do you participate in science outreach activities? = I do not participate in science outreach activities*

For the next series of questions, please indicate if the statement is true, false, or not applicable:

| | True | False | Not Applicable |
|---|---|---|---|
| I am not interested in participating in science outreach | ○ | ○ | ○ |
| I do not have time to participate in science outreach | ○ | ○ | ○ |
| I do not have funding to participate in science outreach | ○ | ○ | ○ |
| My supervisor(s) would not approve of my participation in science outreach | ○ | ○ | ○ |

*Display This Question:*

*If How often do you participate in science outreach activities? = I do not participate in science outreach activities*

You indicated that you do not participate in science outreach. Would you be willing to answer the following statement, "I do not participate in science outreach because..."?

○ No

○ Yes ________________________________________________

*Display This Question:*

*If How often do you participate in science outreach activities? != I do not participate in science outreach activities*

For the next series of questions, please indicate if the statement is true, false, or not applicable: **I participate in science outreach...**

| | True | False | Not Applicable |
|---|---|---|---|
| because it is fun and enjoyable | ○ | ○ | ○ |
| to share my research findings with non-scientists | ○ | ○ | ○ |
| for professional development | ○ | ○ | ○ |
| because of grant/funding requirements | ○ | ○ | ○ |
| for socio-cultural or political reasons | ○ | ○ | ○ |
| to serve as a role model or mentor | ○ | ○ | ○ |
| to improve diversity, equity, and access to STEM | ○ | ○ | ○ |

Page Break

*Display This Question:*

*If How often do you participate in science outreach activities? != I do not participate in science outreach activities*

What are some of the challenges that you face when trying to conduct or participate in science outreach activites? Please select all that apply:

- ▢ Lack of time
- ▢ Lack of funding
- ▢ Lack of support from my peers
- ▢ Lack of support from my department
- ▢ Lack of support from my supervisors
- ▢ Lack of other essential resources (ex. equipment, venue, transportation)
- ▢ Lack of partners or collaborators (ex. school districts, teachers, museums, organizations)
- ▢ Public disinterest interest science
- ▢ Communication or language barriers
- ▢ Other (please specify) ________________________________________________

Page Break

*Display This Question:*

*If How often do you participate in science outreach activities? != I do not participate in science outreach activities*

*Optional Question:*

I participate in science outreach because... [Fill in the blank]

________________________________________________________________

**End of Block: Science Outreach Motivations / Limitations**

**Start of Block: Faculty / PI**

Are you aware of outreach activities or initiatives at your institution?

- Yes
- No
- Unsure
- Not Applicable

Are science outreach activities valued by your tenure and promotion committee?

- Yes
- No
- Unsure
- Not Applicable

Do you think science outreach activities should be valued by tenure and promotion committees?

- Yes
- No
- Unsure
- Not applicable

Does your institution have a dedicated staff, office, or department which coordinates science outreach activities?

- Yes
- No
- Unsure
- Not Applicable

Page Break

Do you support graduate student or post-doctoral fellows' participation in science outreach activities?

- Yes
- No
- Unsure
- Not Applicable

*Display This Question:*

*If Do you support graduate student or post-doctoral fellows' participation in science outreach activ... = No*

You indicated that you do not support graduate student or post-doctoral fellows' participation in science outreach. Would you be willing to provide us with your thoughts on the topic?

- ◯ No
- ◯ Yes ________________________________________________

Do you think science outreach-focused professions are valid career options for STEM trainees?

- ◯ Yes
- ◯ No
- ◯ Unsure
- ◯ Not Applicable

Page Break

*Display This Question:*

*If Does your institution have a dedicated staff, office, or department which coordinates science out... = Yes*

You indicated that your institution has a dedicated staff, office, or department which coordinates science outreach activities.

The science outreach staff, office, or department at my institution:

| | Yes | No | Unsure | Not Applicable |
|---|---|---|---|---|
| Is effective at conducting or supporting science outreach activities or initiatives | ○ | ○ | ○ | ○ |
| Has a positive impact on institutional culture | ○ | ○ | ○ | ○ |
| Has a positive impact on relationships with local community members | ○ | ○ | ○ | ○ |
| Has a positive impact on donor or alumni relations | ○ | ○ | ○ | ○ |

*Display This Question:*

*If Does your institution have a dedicated staff, office, or department which coordinates science out... = Yes*

I have collaborated with the science outreach staff, office, or department at my institution:

○ Yes

○ No

○ Unsure

○ Not Applicable

Page Break

Please select the *primary* source of your research funding:

- ○ National Science Foundation
- ○ National Institutes of Health
- ○ Private foundations or organizations
- ○ Institutional or departmental funds
- ○ Philanthropic funding

Do you support or oppose broader impact requirements for research funding mechanisms?

- ○ Strongly oppose
- ○ Somewhat oppose
- ○ Neutral
- ○ Somewhat favor
- ○ Strongly favor
- ○ I am unfamiliar with broader impact requirements

**End of Block: Faculty / PI**

**Start of Block: Science Outreach Professionals**

You indicated that science outreach is a part of your job description or that you consider yourself to be a science outreach professional. Is this correct?

- ○ Yes
- ○ No

How long have you been employed as science outreach professional?

- Less than 1 year
- 1-2 years
- 2-5 years
- 5-9 years
- 10 or more years

Are you considered a full time or part time employee?

- Full time
- Part time
- Unsure
- Other

Please select your salary range:

- Less than $25,000
- $25,000 to $34,999
- $35,000 to $49,999
- $50,000 to $74,999
- $75,000 to $99,999
- $100,000 to $149,999
- $150,000 or more

On a scale of 1 to 10, how satisfied are you with your current level of compensation?

| | 1 | 2 | 3 | 4 | 5 | 6 | 7 | 8 | 9 | 10 |
|---|---|---|---|---|---|---|---|---|---|---|
| Level of satisfaction (1 being not satisfied at all, 10 being completely satisfied) | | | | | | | | | | |

Page Break

How do you feel you are valued by the following groups:

| | Not at all valued | Slightly valued | Moderately valued | Very valued | Completely valued | Not Applicable |
|---|---|---|---|---|---|---|
| Trainees at your institution | ○ | ○ | ○ | ○ | ○ | ○ |
| Colleagues at your institution | ○ | ○ | ○ | ○ | ○ | ○ |
| Faculty members at your institution | ○ | ○ | ○ | ○ | ○ | ○ |
| Leadership at your institution | ○ | ○ | ○ | ○ | ○ | ○ |
| The broader scientific community | ○ | ○ | ○ | ○ | ○ | ○ |

How easy or difficult is it to obtain the resources that you need to conduct science outreach at your institution?

- Extremely easy
- Somewhat easy
- Neither easy nor difficult
- Somewhat difficult
- Extremely difficult

How easy or difficult was it for you to find employment as a science outreach professional?

- Extremely easy
- Somewhat easy
- Neither easy nor difficult
- Somewhat difficult
- Extremely difficult

In your current role as a science outreach professional, do you view yourself as a scientist?

- Yes
- No
- Unsure
- Not applicable

Page Break

*Display This Question:*

*If In your current role as a science outreach professional, do you view yourself as a scientist?  = No*

You indicated that you do not view yourself as scientist. Would you be willing to share why?

- ◯ No
- ◯ Yes ________________________________________________

*The next question is optional but we would greatly appreciate your response:*
Why did you decide to pursue a career in science outreach -or- how did you end up in your current role as a science outreach professional?

________________________________________________________________

________________________________________________________________

________________________________________________________________

________________________________________________________________

________________________________________________________________

**End of Block: Science Outreach Professionals**

**Start of Block: Leadership**

You indicated that you hold a leadership position or role within your institution. Is this correct?

- ◯ Yes
- ◯ No

Please select your agreement with the following statements:

| | Strongly disagree | Somewhat disagree | Neither agree nor disagree | Somewhat agree | Strongly agree | Not Applicable |
|---|---|---|---|---|---|---|
| I support science outreach activities or initiatives at my institution | ○ | ○ | ○ | ○ | ○ | ○ |
| I think that institutional support for dedicated science outreach personnel is a priority | ○ | ○ | ○ | ○ | ○ | ○ |
| I think that institutional support for a dedicated science outreach office is a priority | ○ | ○ | ○ | ○ | ○ | ○ |
| I have made direct efforts to establish a centralized science outreach program at my institution | ○ | ○ | ○ | ○ | ○ | ○ |

**End of Block: Leadership**
